# QOBRA: A Quantum Operator-Based Autoencoder for *De Novo* Molecular Design

**DOI:** 10.1101/2025.07.15.665010

**Authors:** Yue Yu, Francesco Calcagno, Haote Li, Victor S. Batista

## Abstract

We introduce a variational quantum autoencoder tailored for *de novo* molecular design named QOBRA (Quantum Operator-Based Real-Amplitude autoencoder). QOBRA le circuits for real-amplitude encoding and the SWAP test to estimate reconstruction and latent-space regularization errors during back-propagation. Adjoint encoderand decoder unitary transformations and a generative process that ensures accurate reconstruction as well as novelty, uniqueness, and validity of the generated samples. We showcase QOBRA as applied to *de novo* design of Ca^2+^-, Mg^2+^-, and Zn^2+^-binding metalloproteins after training the generative model with a modest dataset.

**Significance Statement:** Recent advancements in classical generative machine learning have shown significant strides in molecular design for targeted applications. *N*onetheless, these advancements are fundamentally limited by classical computation based on binary units. We introduce a quantum computation-based ML framework employing qubits, which exhibits the ability to synthesize *de novo* molecular instances with specified properties from limited datasets. Quantum networks require exponentially fewer parameters than classical ones, enhancing their trainability and efficiency. While our demonstration focuses on metalloprotein primary sequences, the paradigm is adaptable to diverse molecular designs. This integration of AI and quantum computing holds potential to expand the scientific and technological frontiers of both domains within a practical framework.

---

The design of molecular compounds for targeted functions and applications has long been a cornerstone of chemical research (1, 2). With the rise of computational methods, computer-aided molecular design (CAMD) has advanced significantly, though it continues to face key challenges (3, 4). Early efforts on leveraging structure–function relationships (5, 6) enabled applications ranging from drug delivery to materials science. However, CAMD has remained quite limited due to the complexity of correlating molecular structure with molecular properties in the vast chemical space with a combinatorial number of possible molecules (7, 8).

In recent years, deep learning has driven a new wave of algorithms for molecular design (9, 10). *N*eural networks (*NN*s) can now extract complex, hidden patterns from datasets of lead compounds, enabling the generation of novel molecules with structures and properties informed by those of the training set. In fact, popular AI libraries (e.g., DeepChem (11)) are routinely used to predict molecular properties directly from structure. On the generative side, architectures such as generative adversarial networks (GA*N*s) (12) and reinforcement learning (RL) frameworks (13) can achieve excellent performance for the generation of valid molecules.

Specifically, deep learning models have focused on protein design (9, 14). Proteins are fundamental to life, carrying out a wide range of functions including catalysis (15), transport (16), signaling (17), and regulation (18). They are also implicated in numerous human diseases such as cancer (19), diabetes (20), and Alzheimer’s disease (21), making protein engineering a central challenge in biochemistry. De novo design of proteins thus holds promise for advances in a wide range of applications, including targeted interventions in personalized medicine (22). It has been shown that neural networks can uncover hidden patterns in natural protein sequences and structures, enabling the generation of artificial proteins with enhanced properties and biologically plausible architectures (23, 24). To date, most models have focused on modifying or improving existing protein scaffolds (22), while the space of fully de novo protein design remains comparatively much less explored (23). Greener et al. (25) have reported one application of a classical variational autoencoder (VAE) for protein generation, capable of producing novel peptide sequences that bind metal ions by modifying input sequences of up to 140 amino acids.

Despite recent advances, classical machine learning models remain constrained by rather demanding computational encoding schemes, large neural networks, extensive training data requirements, and significant memory demands. These limitations hinder their scalability and efficient retuning for broader applicability. Quantum machine learning (QML) promises an alternative, introducing a paradigm shift in computation by leveraging variational quantum circuits that can be trained with back-propagation (24). QML models can exploit the encoding efficiency of quantum superposition states with intrinsic parallelism, potentially providing significant efficiency gains.

Superposition states and quantum entanglement should offer key advantages since they can enable the encoding of correlations that are fundamentally unattainable in classical systems (26, 27). Quantum machine learning (QML) models have also demonstrated improved generalization performance and reduced data requirements compared to classical models (28). Moreover, quantum systems can efficiently represent and manipulate exponentially large state spaces. An *N*-qubit system encodes 2^*N*^ states in parallel; for example, 10 qubits represent 1024 states, while 266 qubits represent approximately 10^80^ states—comparable to the number of atoms in the observable universe (29). This combination of exponentially scalable state representation and lower data demands positions QML as a promising approach for domains such as molecular design, where combinatorial complexity and limited training data present major bottlenecks.

Quantum variational autoencoders (QVAEs) are emerging as powerful tools for processing quantum data and simulating quantum systems. These models combine classical variational autoencoders with quantum components to enable efficient compression, representation learning, and generation of quantum states (30, 31). QVAEs have demonstrated competitive performance on tasks like image generation and can be trained using quantum Monte Carlo simulations (30). Recent advancements include the ζ-QVAE, which utilizes regularized mixed-state latent representations and can be applied directly to quantum data (31). Additionally, quantum circuit autoencoders have been developed to compress information within quantum circuits, with applications in anomaly detection and noise mitigation (32). These quantum autoencoder models show promise in learning efficient representations of quantum states, including those that are difficult to simulate classically, suggesting potential applications in near-term quantum hardware (33).

Here, we introduce a QVAE tailored for *de novo* molecular design named **QOBRA** (Quantum Operator-Based Real-Amplitude autoencoder), schematically illustrated in Fig. 1IB. QOBRA is a generative model that learns to encode input data into a continuous, low-dimensional latent space and decode it to reconstruct the original data. Unlike conventional autoencoders, VAEs impose a probabilistic structure—typically a multivariate Gaussian—on the latent space. This regularization enables smooth interpolation between latent representations and conditional generation of molecules in close chemical proximity to a reference structure (10, 25, 34). When appropriately trained, VAEs can generate novel compounds that preserve key characteristics of the training distribution. Prior work has demonstrated their utility across a range of molecular design tasks, including the generation of molecules with tailored physico-chemical properties, selective binding affinities, or compatibility with specific retrosynthetic routes (10, 34, 35), as well as applications in protein design (23, 25) and molecular structure prediction (12, 36). QOBRA is agnostic of the specific quantum computing platform, so we describe how to implement it on conventional qubit-based devices (Part I) as well as on hybrid qubit-qumode platforms (37).

**Fig. 1.**
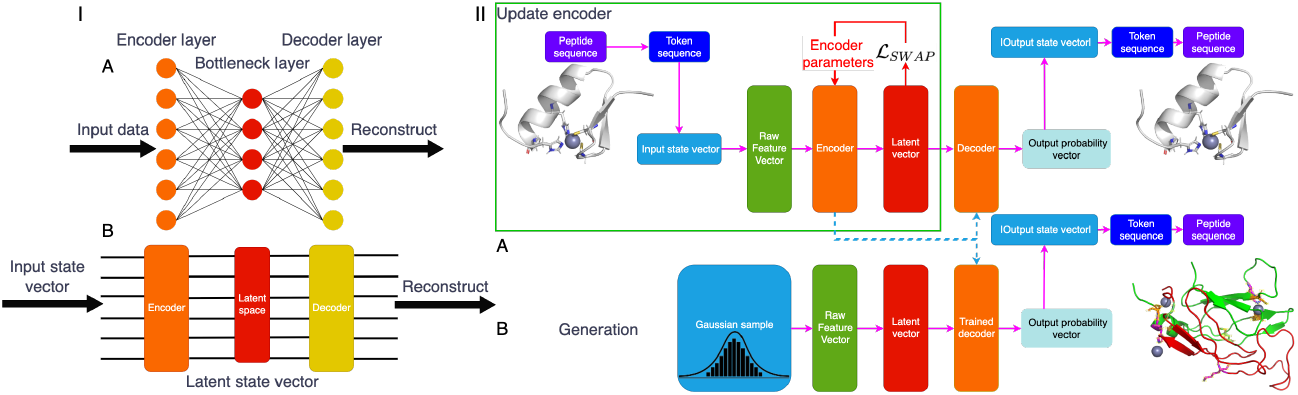
Panel I: Schematic comparison of classical (A) and quantum (B) variational autoencoders. Both architectures include an encoder (orange), a latent space (red), and a decoder (yellow). Panel II: Overview of the QOBRA model. (A) During training, input peptide sequences are embedded into a quantum circuit (encoder), mapped to a latent space, and reconstructed via the decoder—defined as the adjoint of the encoder. (B) After training, new peptide sequences can be generated by sampling from the learned latent space.

In this work, we illustrate QOBRA as applied to *de novo* protein design. Hence, we demonstrate the effectiveness of QOBRA in generating metalloproteins that selectively bind divalent metal ions, including Ca^2+^, Mg^2+^, and Zn^2+^. The model reliably produces appropriate metal-binding sites, as defined by both the primary amino acid sequence and the spatial arrangement of coordinating side chains. QOBRA exhibits strong robustness to hyperparameter variation and consistently delivers high-quality designs using minimal training data and a compact set of variational parameters.

## Results & Discussion

This section presents the results of QOBRA-driven *de novo* generation of Ca^2+^-, Mg^2+^-, and Zn^2+^-binding proteins. We begin by analyzing the impact of key hyperparameters on generation performance, with particular focus on the ansatz unit repetition number (*r*) and the number of qubits (*N*_*q*_). Their influence on both model efficiency and structural quality is systematically investigated.

We then highlight representative metalloproteins generated using the optimal hyperparameter configurations, demonstrating that QOBRA not only produces high-quality protein structures but also outperforms its classical counterpart in key performance metrics.

Generation quality is assessed by comparing features of the generated proteins against those in the training set. Specifically, we examine token frequency distributions, peptide length distributions, the number of ion binding sites, and the number of chains per complex. To quantify the alignment between generated and training data, we compute a relative ratio (*RR*) for each of these four properties. An ideal model would yield *RR* = 1 across all metrics.

In addition, we evaluate the generated sequences using the NUVR metric, which assesses novelty (N), uniqueness (U), validity (V), and reconstruction accuracy (R). Each component is scored between 0 and 1, with 1 indicating a sequence that is entirely novel, unique, chemically valid, and accurately reconstructed. Further methodological details are provided in the Supporting Information (34).

### A. Effect of Ansatz Depth (r**) on Model Performance**

In classical convolutional neural networks, model capacity is strongly influenced by both depth and the number of trainable parameters (38). Analogously, in quantum machine learning (QML), circuit depth plays a critical role in model expressivity and learning performance. In QOBRA, this depth is governed by the number of repetitions *r* of the RA ansatz.

Fig. 2 shows the latent space fitting quality after training QOBRA with *r* = 1 to 3. The inset highlights the first component of the latent vectors, illustrating how the “head” of the sequence is embedded in latent space. The main plots display the fitting behavior of the remaining components. The target latent distribution is a Gaussian with zero mean and standard deviation 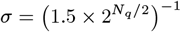.

**Fig. 2.**
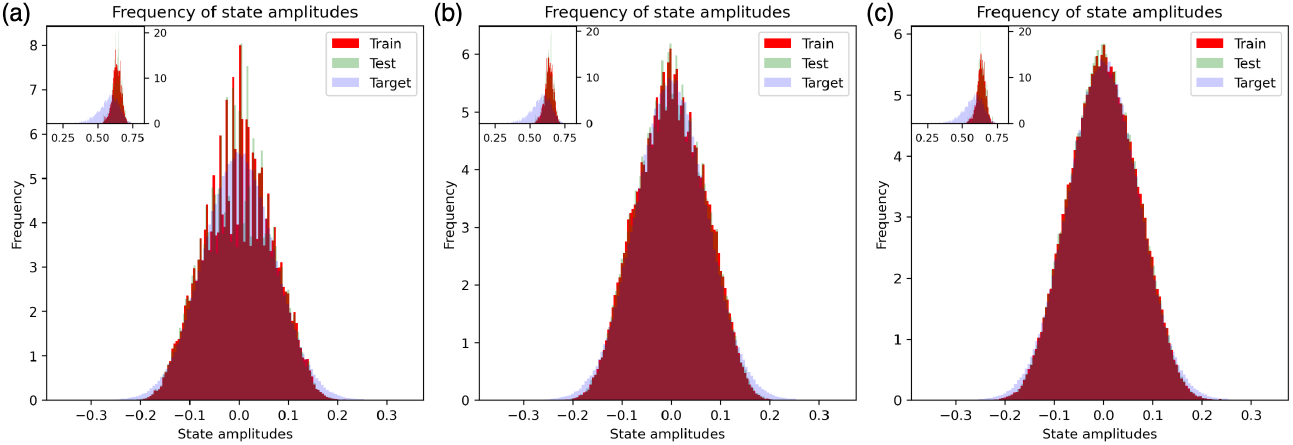
Latent space fitting after training on Zn^2+^ data with *N*_*q*_ = 7 for different ansatz depths: *r* = 1 (a), *r* = 2 (b), and *r* = 3 (c). Increased depth leads to improved alignment with the target distribution, reflecting higher model expressivity.

For *r* = 1 (Fig. 2a), the model shows limited ability to match the target distribution. Increasing to *r* = 2 (Fig. 2b) significantly improves the fit, indicating that a deeper ansatz enhances learning capacity. A further increase to *r* = 3 (Fig. 2c) offers only marginal improvements, suggesting that additional depth yields diminishing returns.

As detailed in Tab. 1, increasing *r* leads to a linear growth in the number of trainable parameters and a corresponding increase in training time. Based on this trade-off between performance and efficiency, we fix *r* = 2 for all subsequent experiments.

**Table 1.** Encoder parameter count and training runtime for Zn^2+^ data as a function of ansatz depth *r*, with *N*_*q*_ = 7. Training was performed using 48 x86 64 Intel CPUs. Only encoder parameters are reported.

| $r$ | Parameters | Training Runtime/h |
| --- | --- | --- |
| 1 | 14 | 3.94 |
| 2 | 21 | 5.90 |
| 3 | 28 | 7.03 |

### B. Trade-off Between Qubit Count (*N*_*q*_ **) and Model Capacity**

Another key hyperparameter is the total number of qubits, *N*_*q*_, which defines the maximum peptide length that the model can process. Specifically, a network with *N*_*q*_ qubits can handle sequences of up to 2^*Nq*^ − 1 residues. If *N*_*q*_ is too small, the model cannot generate sufficiently long or complex sequences to represent functional proteins. On the other hand, while the theoretical advantage of quantum machine learning partly stems from scaling with qubit number (39), increasing *N*_*q*_ leads to a linear growth in the number of trainable parameters. This significantly increases the computational cost and training time. To balance expressivity and efficiency, we restrict our exploration to *N*_*q*_ = 6, 7, 8, and 9, as shown in Tabs. 2 and 3.

While the *N*UVR metric remains relatively consistent across the three ion datasets (Tab. 2), the relative ratio (RR) results—summarized in Tab. 3—highlight a more nuanced dependence on the qubit count *N*_*q*_. In general, performance improves with increasing *N*_*q*_, as reflected by RR values approaching the ideal value of 1 across all training scenarios. This trend is most pronounced for Zn^2+^ at *N*_*q*_ = 9, as shown in Fig. 3, where the generated sequences closely match the training distribution across all evaluated metrics: token frequency, number of chains, complex size, and number of binding sites. A broader analysis across all three ions reinforces this pattern. For both Ca^2+^ and Zn^2+^, the *RR* values consistently converge toward 1 as *N*_*q*_ increases—ranging from 0.51 ± 0.68 to 5.05 ± 11.28 for Ca^2+^, and from 1.01 ± 1.39 to 4.52 ± 5.72 for Zn^2+^. Although Mg^2+^ exhibits greater variance and less favorable alignment with the training distribution (RR range: 2.58 ± 2.20 to 10.07 ± 26.03), the underlying trend of improved distributional similarity with increasing *N*_*q*_ remains consistent. Based on this observation, we fix *N*_*q*_ = 9 in all subsequent experiments, enabling the model to generate primary sequences of up to 511 amino acids.

**Table 2.**
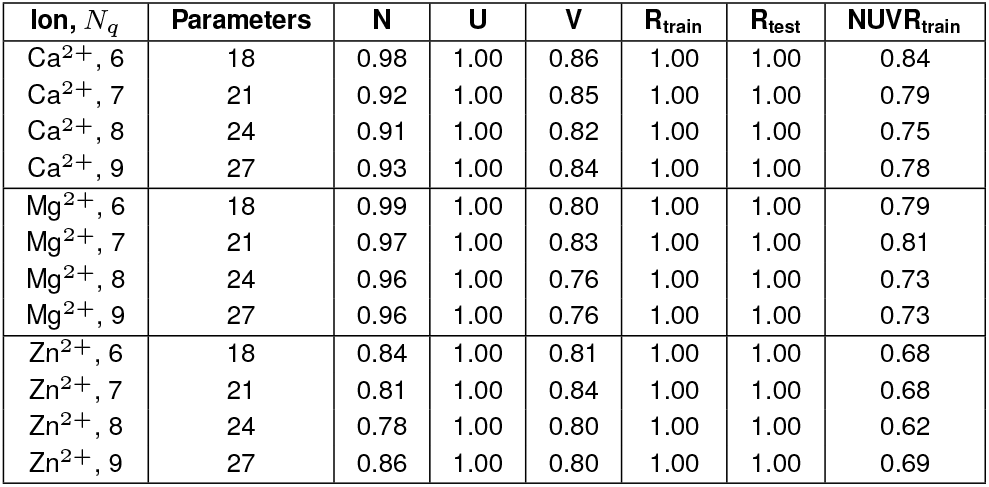
*N*UVR metric components—novelty (*N*), uniqueness (U), validity (V), and reconstruction accuracy (*R*)—for generated sequences, evaluated on training and test sets. Results are shown for each ion type and qubit count *N*_*q*_. The composite *N*UVR_train_ score reflects generation quality under each configuration.

| Ion, $N_q$ | Parameters | N | U | V | $R_{\text{train}}$ | $R_{\text{test}}$ | NUVR<sub>train</sub> |
| --- | --- | --- | --- | --- | --- | --- | --- |
| $\text{Ca}^{2+}$, 6 | 18 | 0.98 | 1.00 | 0.86 | 1.00 | 1.00 | 0.84 |
| $\text{Ca}^{2+}$, 7 | 21 | 0.92 | 1.00 | 0.85 | 1.00 | 1.00 | 0.79 |
| $\text{Ca}^{2+}$, 8 | 24 | 0.91 | 1.00 | 0.82 | 1.00 | 1.00 | 0.75 |
| $\text{Ca}^{2+}$, 9 | 27 | 0.93 | 1.00 | 0.84 | 1.00 | 1.00 | 0.78 |
| $\text{Mg}^{2+}$, 6 | 18 | 0.99 | 1.00 | 0.80 | 1.00 | 1.00 | 0.79 |
| $\text{Mg}^{2+}$, 7 | 21 | 0.97 | 1.00 | 0.83 | 1.00 | 1.00 | 0.81 |
| $\text{Mg}^{2+}$, 8 | 24 | 0.96 | 1.00 | 0.76 | 1.00 | 1.00 | 0.73 |
| $\text{Mg}^{2+}$, 9 | 27 | 0.96 | 1.00 | 0.76 | 1.00 | 1.00 | 0.73 |
| $\text{Zn}^{2+}$, 6 | 18 | 0.84 | 1.00 | 0.81 | 1.00 | 1.00 | 0.68 |
| $\text{Zn}^{2+}$, 7 | 21 | 0.81 | 1.00 | 0.84 | 1.00 | 1.00 | 0.68 |
| $\text{Zn}^{2+}$, 8 | 24 | 0.78 | 1.00 | 0.80 | 1.00 | 1.00 | 0.62 |
| $\text{Zn}^{2+}$, 9 | 27 | 0.86 | 1.00 | 0.80 | 1.00 | 1.00 | 0.69 |

**Table 3.** Relative ratio (RR) metrics for token frequency, number of chains, peptide length, and binding sites, computed across different ion types and qubit counts (*N*_*q*_). Each row also lists the total number of encoder parameters. Higher *N*_*q*_ allows longer sequences but increases model complexity.

| Ion, $N_q$ | Parameters | Token Freq. | Chains | Length | Binding Sites |
| --- | --- | --- | --- | --- | --- |
| $\text{Ca}^{2+}$, 6 | 18 | $1.69 \pm 2.01$ | $2.75 \pm 3.42$ | $9.85 \pm 8.21$ | $4.89 \pm 8.35$ |
| $\text{Ca}^{2+}$, 7 | 21 | $2.13 \pm 2.23$ | $7.62 \pm 8.58$ | $23.71 \pm 27.47$ | $3.24 \pm 3.85$ |
| $\text{Ca}^{2+}$, 8 | 24 | $1.82 \pm 1.31$ | $9.91 \pm 13.06$ | $13.34 \pm 23.09$ | $1.12 \pm 0.94$ |
| $\text{Ca}^{2+}$, 9 | 27 | $1.12 \pm 0.62$ | $1.05 \pm 0.60$ | $5.05 \pm 11.28$ | $0.51 \pm 0.68$ |
| $\text{Mg}^{2+}$, 6 | 18 | $4.15 \pm 9.20$ | $16.34 \pm 14.80$ | $23.12 \pm 22.18$ | $27.35 \pm 37.99$ |
| $\text{Mg}^{2+}$, 7 | 21 | $6.06 \pm 12.01$ | $30.95 \pm 36.58$ | $46.94 \pm 47.87$ | $46.26 \pm 47.54$ |
| $\text{Mg}^{2+}$, 8 | 24 | $5.04 \pm 5.77$ | $30.38 \pm 48.18$ | $23.09 \pm 43.33$ | $21.71 \pm 26.31$ |
| $\text{Mg}^{2+}$, 9 | 27 | $2.58 \pm 2.20$ | $4.07 \pm 4.01$ | $10.07 \pm 26.03$ | $4.55 \pm 4.04$ |
| $\text{Zn}^{2+}$, 6 | 18 | $14.43 \pm 40.71$ | $2.37 \pm 2.91$ | $3.75 \pm 2.13$ | $7.72 \pm 11.48$ |
| $\text{Zn}^{2+}$, 7 | 21 | $17.36 \pm 40.76$ | $3.40 \pm 5.23$ | $4.24 \pm 1.89$ | $5.25 \pm 5.22$ |
| $\text{Zn}^{2+}$, 8 | 24 | $11.35 \pm 14.92$ | $2.68 \pm 5.40$ | $3.36 \pm 2.26$ | $3.38 \pm 4.31$ |
| $\text{Zn}^{2+}$, 9 | 27 | $4.52 \pm 5.72$ | $1.19 \pm 1.51$ | $1.73 \pm 1.37$ | $1.01 \pm 1.39$ |

**Fig. 3.**
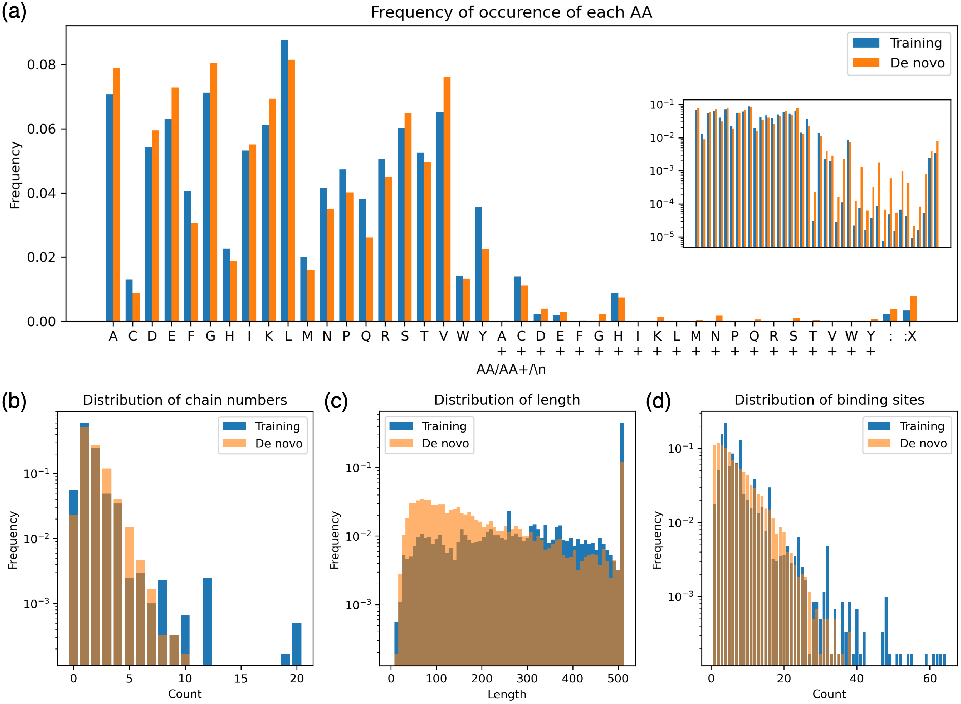
Histograms for Zn^2+^ with *N*_*q*_ = 9 of the frequencies of tokens (a), chain numbers (b), peptide lengths (c), and ion binding sites (d) comparing generated sequences (orange) to the training set (blue). The length is calculated as the number of AAs plus : in a sequence, while chain number is computed as how many : a sequence contains. A 0 chain number implies that the sequence is a partial domain within a larger complex. In (a), the inset plot shows the same as the main plot, but with a log-scale y-axis.

### C. Tertiary Structure Prediction and Refinement

In *de novo* metalloprotein design, accurate reconstruction of tertiary structure from a generated primary sequence is essential for assessing functional viability—particularly for identifying and localizing metal ion binding sites. To enable this, we implemented a structure prediction pipeline tailored to QOBRA-generated sequences (Fig. 4).

**Fig. 4.**
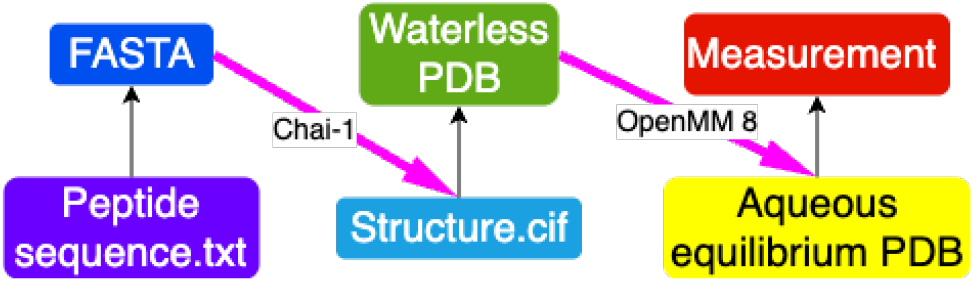
Schematic overview of the sequence-to-structure pipeline. A generated primary sequence is formatted and converted to FASTA, processed by the Chai-1 language model to predict a three-dimensional structure (in CIF and PDB formats), and subsequently equilibrated via molecular dynamics simulation in OpenMM 8 to produce a solvated, biologically relevant structure.

#### C.1 Sequence-to-Structure Workflow

Fig. 4 illustrates the computational pipeline used to convert a generated primary sequence into a fully solvated, structurally equilibrated protein model suitable for downstream analysis. The workflow consists of four main stages:

1. **Sequence Input and Formatting:** The pipeline begins with a peptide sequence generated by QOBRA, stored in a plain text file (**sequence.txt**). This sequence is converted into a standard **FASTA** format to ensure compatibility with structure prediction tools.
2. **Structure Prediction (Chai-1):** The FASTA file is processed by the Chai-1 structure prediction engine (40), which outputs a predicted 3D conformation in .cif format. This file is then converted to a PDB format representing the protein atomic coordinates in the absence of solvent and ions—referred to as the *dry PDB*.
3. **Solvation and Molecular Dynamics Simulation (OpenMM 8):** The dry structure is input into OpenMM 8 (41), where it is solvated using the TIP3P water model and neutralized with counterions. A molecular dynamics (MD) simulation is then performed to equilibrate the structure under near-physiological conditions. The resulting output is an equilibrated *aqueous PDB* that incorporates solvent and ion coordination effects.
4. **Structural Analysis:** The equilibrated structure is subsequently subjected to structural analysis, including RMSD calculations and evaluation of binding site integrity. These measurements provide insight into the physical plausibility and stability of the *de novo* generated protein models.

This modular workflow enables reliable translation of synthetic sequences into realistic 3D structures for functional and biophysical characterization.

#### C.2 Three-Dimensional Protein Structures

Three-dimensional structure prediction was performed using Chai-1 (40), a state-of-the-art deep learning framework for modeling protein conformations. Representative outputs of Chai-1 applied to QOBRA-generated sequences are shown in Fig. 5. This task presents a nontrivial challenge: the generated sequences are synthetic and lack homologs in structural databases, precluding the use of homology-based modeling. Consequently, Chai-1 infers structural configurations in a purely *ab initio* manner. Metal ion placement is handled iteratively, with ions introduced into the structure until all predicted coordination sites are saturated based on local residue geometry.

**Fig. 5.**
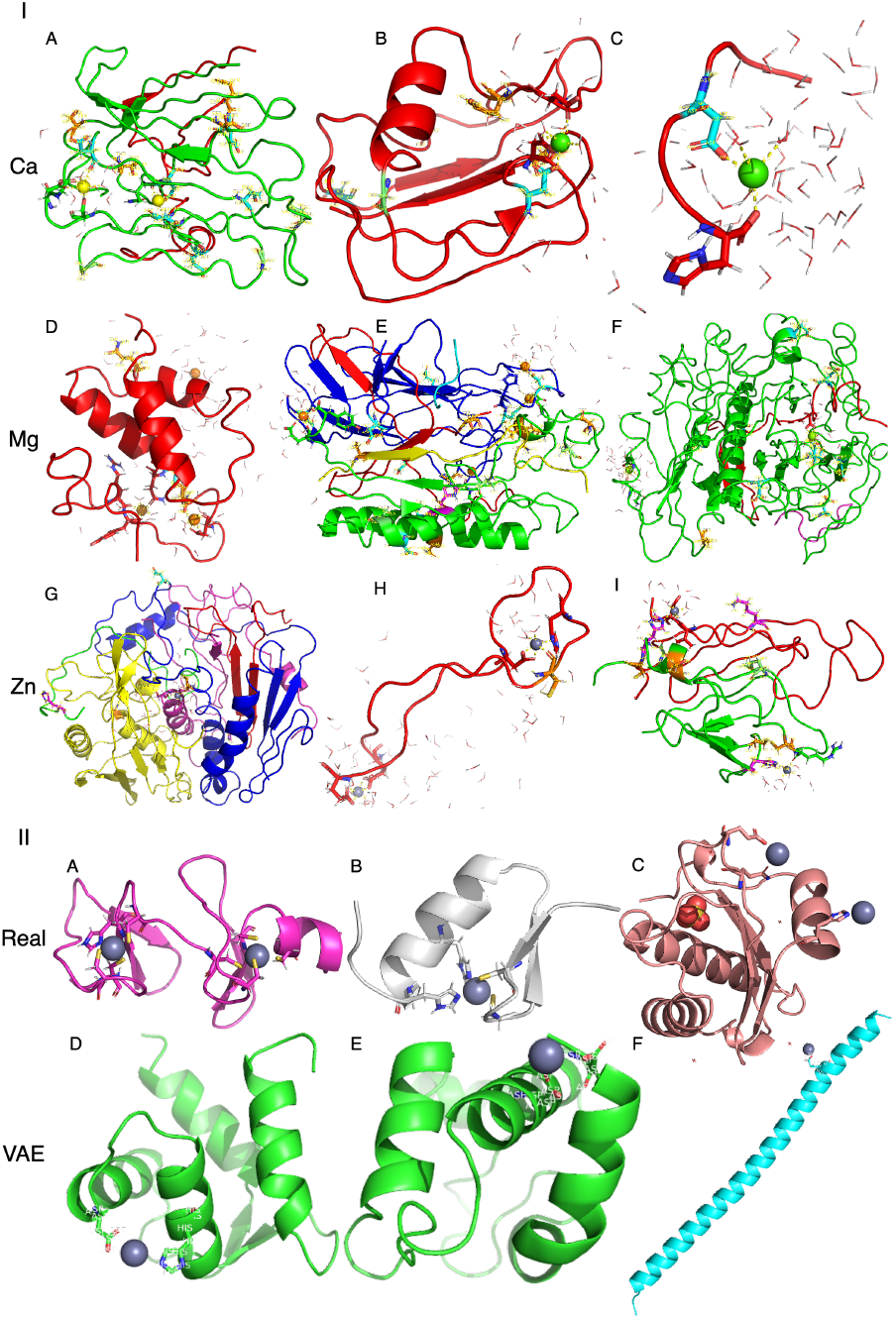
(A) Representative artificial metalloproteins generated by QOBRA with *N*_*q*_ = 9 and *r* = 2. Structures include Ca^2+^-binding (green, A1–A3), Mg^2+^-binding (lime, A4–A6), and Zn^2+^-binding (gray, A7–A9) proteins. Tertiary structures were predicted using Chai-1 (40). Highlighted residues indicate predicted ion-coordinating sites identified by the QOBRA model. Coordinating water molecules are also shown, forming metal-specific coordination geometries—hexahedral for Ca^2+^ and Mg^2+^, tetrahedral for Zn^2+^. (B) Examples of Zn^2+^-binding proteins from nature (B1–B3) and from sequences generated by the VAE model of Greener et al. (B4–B6).

To ensure structural and physicochemical plausibility, all predicted conformations were subjected to molecular dynamics (MD) refinement in explicit solvent. Simulations were carried out using OpenMM 8 (41) at a constant temperature of 300 K. Protein interactions were described using the AMBER14 force field (42), while solvent was modeled using the TIP3P water model (43). Each structure was solvated in a cubic water box extending 0.5 nm beyond the protein in all dimensions, and counterions (*N*a^+^, Cl^*−*^) were added to neutralize net charge.

Systems underwent energy minimization using Langevin dynamics for 50,000 steps, followed by temperature equilibration to 300 K via a Langevin thermostat (44), employing a 4 fs integration timestep and a friction coefficient of 1 ps^*−*1^ over an additional 50,000 steps. Structural stability and convergence were assessed throughout the simulation using root-mean-square deviation (RMSD) analysis, calculated with MDTraj (45). This refinement pipeline produces solvent-equilibrated structures, allowing direct comparison to natural metalloproteins and enabling downstream biophysical or functional analysis.

#### C.3 Selectivity and Specificity

The primary sequences generated by QOBRA contain canonical secondary structure elements, including α-helices, β-sheets, and coils—in proportions comparable to those observed in the training set (α-helices: 30–45 %; β-sheets: 20–30 %; loops/turns/other: 25–40 %). These sequences fold into tertiary structures that closely resemble those of natural proteins, as illustrated by representative examples in Fig. 5A. Furthermore, the predicted metal-binding sites agree with established principles of coordination chemistry with preferred ligands of amino acid side chains, distinct for each type of metal.

We define a Chai-1 prediction as successful if the predicted 3D structure places a metal ion in close proximity to the residues identified by QOBRA as metal-coordinating. False positives (FP) occur when predicted coordinating residues lack nearby metal ions, while false negatives (F*N*) are residues not predicted by QOBRA but are located near metal ions in the structure. True positives (TP) and true negatives (T*N*) follow the standard definitions. From these, we compute sensitivity and specificity, as follows:

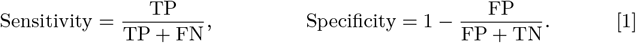

We have evaluated 100 generated structures per metalloprotein type. Coordination was assessed using metal-specific cutoff distances, identifying coordinating atoms from side chains or water molecules. Histogram distributions of sensitivity and specificity are shown in Fig. S3.

Overall, the model achieves high specificity, with most values in the (0.9, 1.0) range across all three ions. Sensitivity, however, is more variable, often peaking near zero, indicating missed coordinating residues. *N*onetheless, occasional cases of 100% sensitivity demonstrate that the model is capable of high performance under the right structural conditions.

Simulations involving Zn^2+^ consistently show coordination pockets composed of residues known to bind Zn^2+^ biologically—histidine, cysteine, aspartate, and glutamate—along with water molecules. These tertiary motifs, consistent with natural and engineered proteins (46, 47), also emerge in QOBRA-derived structures. Similar trends are observed for Ca^2+^ and Mg^2+^, which preferentially coordinate with aspartate, glutamate, and water (48, 49). The predicted binding pockets typically include both QOBRA-predicted residues and additional structural contributors to the coordination sphere.

#### C.4 Coordination Number

A more rigorous assessment of the structural quality of the generated protein models can be obtained by analyzing the coordination number of the bound metal ions—that is, the number of atoms directly coordinating each ion. Coordination numbers are ion-specific and are influenced by both the identity of the ion and the nature of its ligands, including water and non-peptidic molecules (50, 51).

In aqueous protein environments, calcium (Ca^2+^) typically adopts coordination numbers of 6 to 8, magnesium (Mg^2+^) commonly coordinates with 6 atoms, and zinc (Zn^2+^) generally exhibits coordination numbers between 4 and 6. Each ion also has characteristic coordination distances that reflect its size and preferred ligand geometries.

Fig. S4 presents the coordination numbers and corresponding distances observed in our generated structures. *N*otably, the computed cutoff distances required to capture coordinating atoms are consistently shorter than those observed in experimentally determined protein structures. This suggests that the generated proteins may exhibit stronger metal-binding affinities in aqueous environments compared to their natural counterparts, potentially due to tighter coordination geometries.

#### C.5 Secondary Structure Proportions

Table 4 illustrates the proportions of the three secondary structures of proteins for each ion set, with comparison to the expected range for natural proteins. The measurements were performed using DSSP (52) in Biopython (53), which provides a result closely aligning with natural ranges, notwithstanding the tendency of protein language models such as Chai-1 to predict helical structures (40, 54, 55). This suggests that QOBRA possesses a degree of capability to understand protein primary sequence composition to create proxy-natural proportions of domains.

**Table 4.** Protein secondary structure proportions in three types of generated structural sets vs. proportions in natural proteins.

| | α-helix | β-sheet | Coil |
| --- | --- | --- | --- |
| Ca<sup>2+</sup> | 0.30 ± 0.20 | 0.24 ± 0.15 | 0.46 ± 0.14 |
| Mg<sup>2+</sup> | 0.34 ± 0.20 | 0.21 ± 0.15 | 0.45 ± 0.15 |
| Zn<sup>2+</sup> | 0.34 ± 0.20 | 0.20 ± 0.15 | 0.46 ± 0.12 |
| Natural | [0.3, 0.35] | [0.2, 0.25] | [0.4, 0.5] |

#### C.6 Natural vs. Generated

The synergy between the generative capabilities of QOBRA and the structure prediction provided by Chai-1 demonstrates an effective approach for designing protein sequences and structures. Both models recover fundamental biophysical patterns and generate novel proteins that closely replicate the composition and architecture of natural systems. This level of performance is especially notable given the minimal parameter count—only 27 trainable variables—and the modest training set size of approximately 6,000 sequences per metal ion. In comparison, a generative classical model employed a conventional variational autoencoder with 912 neurons, four hidden layers, and more than 105,000 sequences to achieve similar results (Fig. 5B) (25). These outcomes are made possible by the distinctive architecture and operational principles of QOBRA, which differ substantially from those of classical machine learning methods.

## Materials & Methods

An overview of the QOBRA workflow is shown in Fig. 1IIA. The architecture consists of two components: a *quantum encoder* and a *quantum decoder*. Both are implemented as parameterized quantum circuits, mirroring the structure of classical neural networks (c*NN*s). The circuit variational parameters are optimized during training by back-propagation.

In our applications for *de novo* protein design, input peptide sequences are mapped into quantum amplitudes through a letter-to-number encoding scheme, followed by normalization. This transforms discrete token sequences into continuous quantum state vectors suitable for processing by the quantum encoder.

Training jointly optimizes two loss functions in a self-consistent loop, ensuring regularization into a Gaussian latent space distribution and accurate reconstruction through direct comparisons with SWAP tests (detailed in Sec. E).

Following training, the decoder operates independently (Fig. 1IIB) to generate novel peptide sequences. This is achieved by sampling from the latent space, applying the decoder gates to the sampled vectors, and measuring the output quantum state in the computational basis. The resulting probability distribution is square-rooted and mapped back to the closest amino acid token magnitudes, enabling the reconstruction of new peptide sequences.

### D. Encoding Scheme

QOBRA operates on primary amino acid sequences by converting each peptide into a unique real amplitude vector. These vectors are then element-wise square-rooted to produce normalized state vectors, which serve as quantum inputs to the model. All 20 canonical amino acids are represented in the encoding. To differentiate between metal-binding and non-binding residues, two categories are defined: **AA** refers to an amino acid that is not coordinated to a metal ion, while **AA+** designates a metal-binding variant. Each **AA** and **AA+** is assigned a unique numeric token.

Two special tokens are also introduced:

- : – Denotes chain breaks in multi-chain peptides.
- :`X` – Indicates the end of a sequence. Since the number of qubits (*N*_*q*_) determines the dimensionality of the quantum state vector, sequences shorter than 2^*N q*^ − 1 residues are padded by appending : `X`, followed by repeated copies of the peptide. Sequences exceeding this length are truncated at the : `X` marker.

This encoding scheme establishes a consistent and reversible mapping from biological peptide sequences to fixed-dimension quantum state vectors, enabling efficient quantum processing of peptides with diverse lengths and structural features.

Many classical machine learning models are invariant to the absolute values and ordering of input tokens (56, 57). However, the specific choice of token-to-value mapping significantly impacts the model performance of our quantum encoding. This is due to the sensitivity of the circuit to the input vector distribution. For instance, if two tokens with close numerical values—such as free Aspartic acid (`D`) and its ion-bound form (`D+`)—occur at similar frequencies in the training data, the encoder’s intrinsic noise can lead to ambiguity between them. This results in an oversampling of the less frequent token due to value overlap in the latent space. To mitigate this, tokens are assigned numerical values that follow a bell-shaped distribution centered at zero, as shown in Fig. S1. This ensures sufficient separation between tokens, especially for low-frequency ones. Additionally, because quantum measurements return probabilities—i.e., the squared amplitudes—any phase information (sign of the amplitude) is lost. To address this, all token values are assigned unique absolute magnitudes to preserve distinguishability.

Amplitude encoding is normalized, but peptide sequences may vary in length and total token value. To ensure a bijective and decodable representation, we prepend each vector with a fixed constant *n*. This scalar acts as an internal normalization reference, allowing for rescaling and accurate reconstruction of the original sequence. This format ensures compatibility with amplitude encoding while retaining biological interpretability. For example, the peptide sequence `GC · · · LDAE` is mapped, as follows:

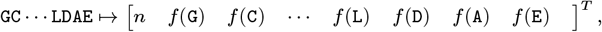

where *f*() assigns a distinct real-valued amplitude to each input token, as defined by a given dictionary.

### E. Loss Functions

Within the autoencoder framework, the model must learn to both encode and decode—that is, to accurately reconstruct—any input sequence from the training set. At the same time, the distribution of encoded vectors in the *latent space* must approximate a well-defined, tractable probability distribution to enable generative sampling (Fig. 1). To enforce reconstructive symmetry, the decoder is implemented as the inverse of the encoder ansatz, specifically as its adjoint operator. This architectural constraint reduces the number of trainable parameters through the optimization of the encoder, ensuring that it maps inputs into a latent representation that supports both reconstruction and generation.

In classical autoencoders, alignment of the latent space with a reference distribution is commonly achieved using the maximum mean discrepancy (MMD) loss (58), or its modified form (m-MMD) (34). This loss encourages the distribution of latent vectors—obtained from the training set—to match a predefined prior, typically a multivariate Gaussian. The m-MMD loss is given by:

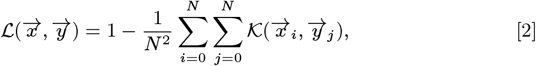

Where 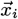 are latent representations from the encoder, and 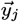 are reference vectors sampled from the prior. The kernel function К is defined as:

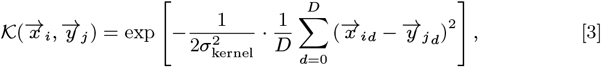

where ***D*** is the dimensionality of the latent space, and σkernel is a tunable bandwidth parameter. This loss promotes statistical alignment between the encoded latent distribution and the reference prior, thereby allowing the decoder to generate meaningful outputs from unseen latent vectors. In practice, all 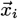 vectors are obtained from the encoder, while the 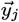 vectors are drawn from the target distribution. To maintain norm consistency, the first element of each 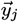 vector encodes a normalization factor, ensuring unit-norm latent states.

A challenge arises when trying to implement this framework on quantum devices since comparing quantum state amplitudes requires quantum state tomography (59) which would not be practical since it is computationally demanding. Here, we bypass the need for quantum state tomography by using the SWAP test (60, 61). As described below, measurements of an ancilla qubit provide an estimate of the overlap between quantum states without having to determine the quantum state amplitudes (Fig. 6I). Accordingly, losses involving vector similarity are reformulated using the SWAP test.

**Fig. 6.**
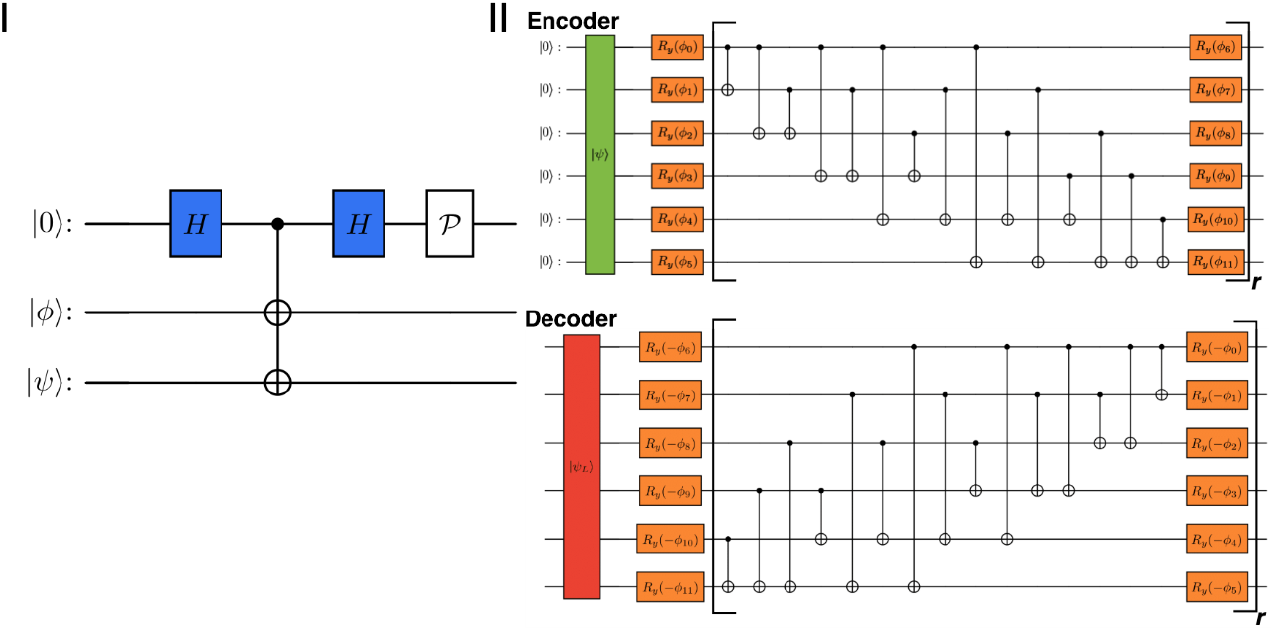
Panel I: Illustration of the SWAP test mechanism. The circuit ends with a probability measurement (P) of the auxiliary qubit on top. When 𝒫 (|0⟩) = 1, the states |*ϕ*⟩ and |*ψ*⟩ are identical. Panel II: 6-qubit RealAmplitutes encoder and decoder ansatz are reported. The state vector input encoding layer (in green for the encoder and in red for the decoder). Rotation gates with trainable parameters are marked in orange. The repetition unit is highlighted with square brackets and the hyperparameter *r*.

Starting with ℒ, defined according to Eq. 2, we redefine the similarity kernel function by using the infidelity, as follows:

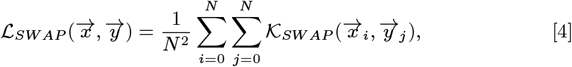

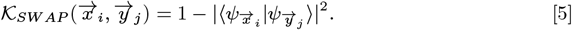

Therefore, the loss implemented by QOBRA is essentially a quantum analogue of the m-MMD loss implemented by the classical kernel-elastic autoencoder (34).

### F. Encoder and Decoder

During training, QOBRA maps the input data into a latent space representation by implementing unitary transformations (24). The architecture of the encoder—i.e., the choice of ansatz—is critical, as it must balance expressiveness with trainability and hardware efficiency. Just as architectural choices define the learning capacity of a classical neural network, the quantum ansatz determines the representational power of the resulting QML model.

For qubit-based platforms, we adopt the fully entangled RealAmplitudes (RA) ansatz (Fig. 6II). As implemented in Qiskit (62), the RA ansatz comprises layers of R_*y*_ rotation gates interleaved with full entanglement operations. All qubits are initialized in the |0⟩ state, and the use of R_*y*_ gates restricts state vectors to the XZ-plane of the Bloch sphere. This ensures real-valued amplitudes, which could permit the use of real-valued loss functions (63).

Fig. 6II illustrates the simplest form of the RA ansatz, with a repetition depth *r* = 1. Higher expressiveness can be achieved by increasing *r*, which adds RA layers with additional trainable parameters. Therefore, the number of trainable parameters in the encoder scales as *N*_*q*_ (*r* + 1), where *N*_*q*_ is the number of qubits. Unless otherwise specified, all results in this study are based on circuits with 6–9 qubits, enabling the representation of proteins with up to 63–511 amino acid residues. The decoder, illustrated in Fig. 6II, is constructed as the complex conjugate (adjoint) of the encoder circuit. It receives the latent vector | ψ_*L*_⟩ as input and shares the same parameter values as the encoder. This architecture ensures efficient and consistent reconstruction in the quantum autoencoding pipeline.

### G. Dataset

We used three curated datasets, publicly available via the QOBRA GitHub repository (64), each corresponding to protein complexes binding a specific divalent metal ion: Ca^2+^, Mg^2+^, or Zn^2+^. The datasets include 13,279 proteins with Ca^2+^-binding, 16,506 with Mg^2+^-binding, and 19,474 proteins binding Zn^2+^. Structures were retrieved from the RCSB Protein Data Bank (PDB) (65) by filtering for entries annotated as containing the respective metal ion.

Because RCSB annotations often conflate structural cofactors with loosely associated ions (66, 67), we applied a post-processing step to remove entries in which the metal ion was not covalently or coordinately bound to amino acid residues. All datasets were partitioned into training and testing sets using a 5:1 ratio.

## ACKNOWLEDGMENTS

V.S.B. acknowledges partial support from Boehringer Ingelheim and computational resources from the *N*ational Energy Research Scientific Computing Center (*N*ERSC), a U.S. Department of Energy Office of Science User Facility located at Lawrence Berkeley *N*ational Laboratory. F.C. acknowledges the “Ing. Luciano Toso Montanari” Foundation for financially supporting his secondment at Yale University (USA) in the group of Professor Victor S. Batista.

## 1. Code and Data Availability

The code and data are available at the QOBRA Github (64). For inquiries, please contact.

