## Supplementary material for "QOBRA: A Quantum Operator-Based Autoencoder for *De Novo* Molecular Design": SI

#### This PDF file includes:

Supporting text
Figs. S1 to S4
SI References

### Supporting Information Text

### Software and Hardware

All computations were executed using Python 3.11.5. The behavioral emulation of a quantum device via classical computation was performed using the Qiskit (version 1.2.0) quantum simulation package. Quantum state measurements were performed using the StateVector simulator (1). Circuit parameter optimization was achieved using the COBYLA optimizer, as implemented within Qiskit. Tensor calculations were facilitated by the PyTorch version 2.2.0 + cu121 package (2). A comprehensive list of libraries used in this study is provided in this link. All simulations delineated in this study were performed utilizing 16 processors in the following hardware configuration: AMD Perlmutter EPYC CPUs equipped with 512 GB of RAM and NVIDIA A100 Tensor Core GPUs, featuring 40 GB of HBM2.

### Encoding scheme

Fig. S1 illustrates the distribution of occurrence frequencies in relation to the square root of token values corresponding to $Zn^{2+}$-protein sequences ranging from 0 to 512 residues in length. As the occurrence frequency increases within the dataset, the corresponding value approaches 0.

### Structures

The assessment of equilibrium is conducted through the measurement of the root-mean-square deviation (RMSD) of the protein complex and its associated metal ions in relation to their initial positions at various intervals throughout the simulation, until stabilization is achieved. Fig. S2 illustrates an example of the RMSD for a protein complex and metal ion relative to their initial positional state throughout the simulation steps within OpenMM 8. The observation of a plateau indicates that the three-dimensional structure has reached equilibrium stabilization.

### Selectivity and Specificity

Fig. S3 presents a histogram illustrating the selectivity and specificity of the predicted coordination residues within selected computationally generated structures, each containing a single ion, across all three ion types.

### Coordination numbers

Comparison of metal coordination number frequency for $Ca^{2+}$-, $Mg^{2+}$-, and $Zn^{2+}$-proteins in the real case (Fig. S4a-c) against generated structures containing 1 (d-f) and 4 (g-i) ions. The cutoffs are chosen so that the majority of coordination numbers match with theoretical values.

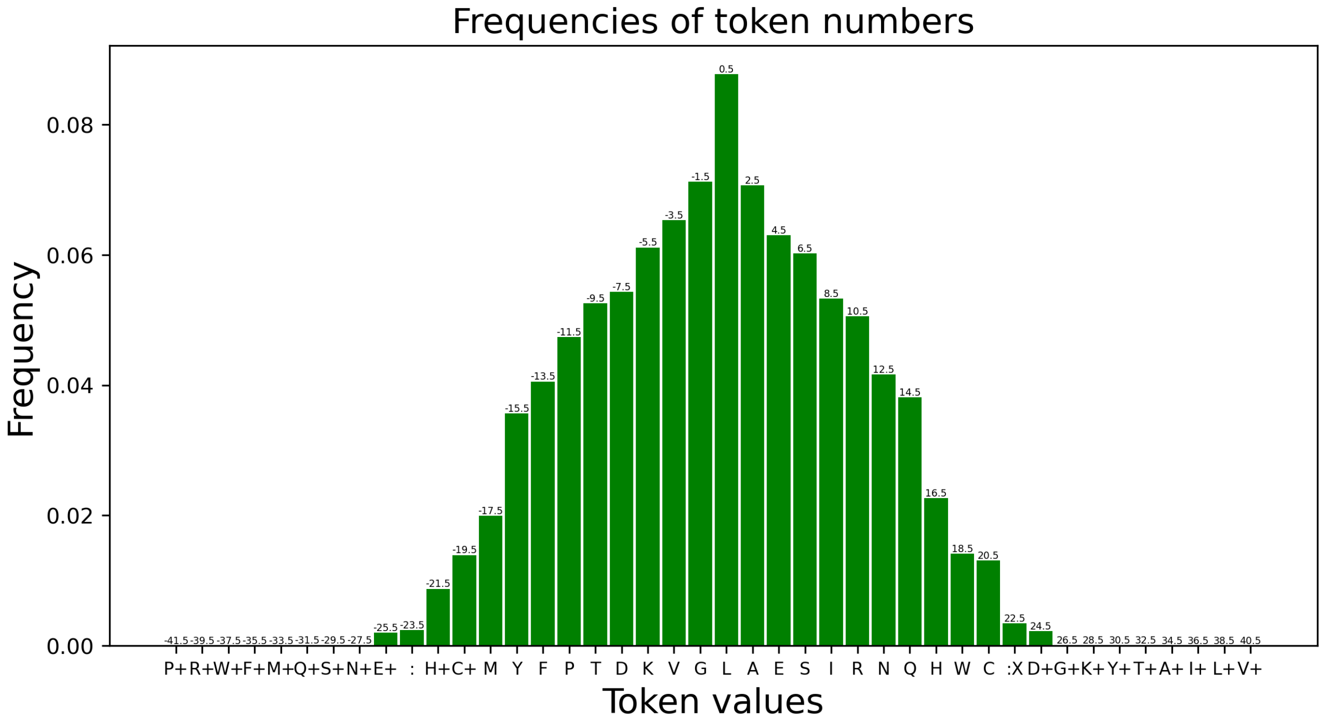
Frequencies of token numbers
Frequency
Token values
0.00
0.02
0.04
0.06
0.08
P+ R+ W+ F+ M+ Q+ S+ N+ E+ : H+ C+ M Y F P T D K V G L A E S I R N Q H W C :X D+ G+ K+ Y+ T+ A+ I+ L+ V+
-41.5 -39.5 -37.5 -35.5 -33.5 -31.5 -29.5 -27.5 -25.5 -23.5 -21.5 -19.5 -17.5 -15.5 -13.5 -11.5 -9.5 -7.5 -5.5 -3.5 -1.5 0.5 2.5 4.5 6.5 8.5 10.5 12.5 14.5 16.5 18.5 20.5 22.5 24.5 26.5 28.5 30.5 32.5 34.5 36.5 38.5 40.5

**Fig. S1**

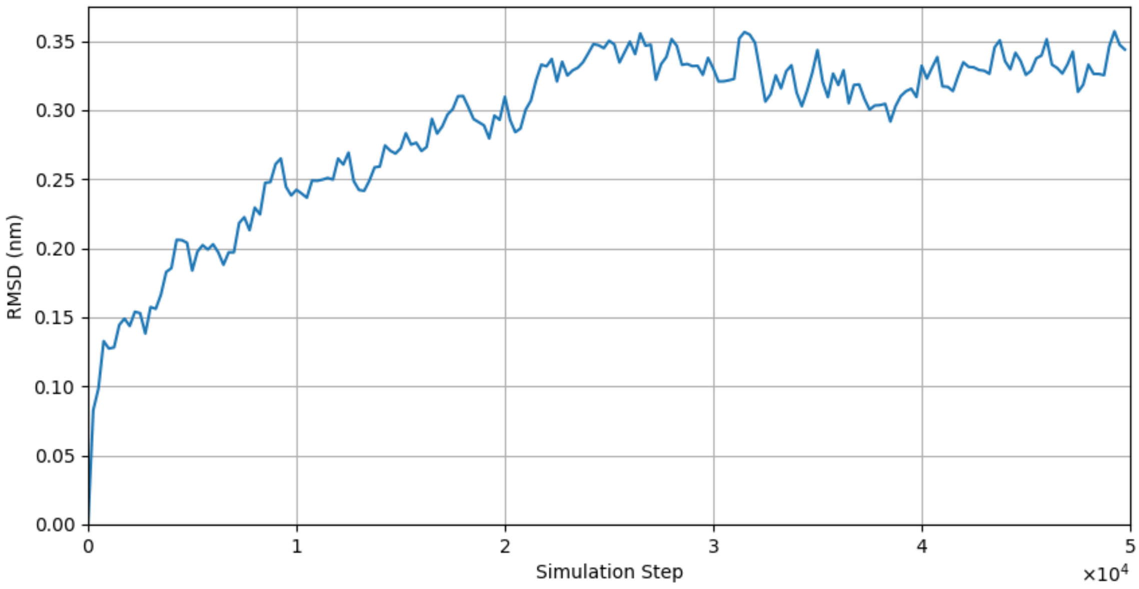

0.35
0.30
0.25
0.20
0.15
0.10
0.05
0.00
RMSD (nm)
0
1
2
3
4
5
Simulation Step
×10^4


**Fig. S2**

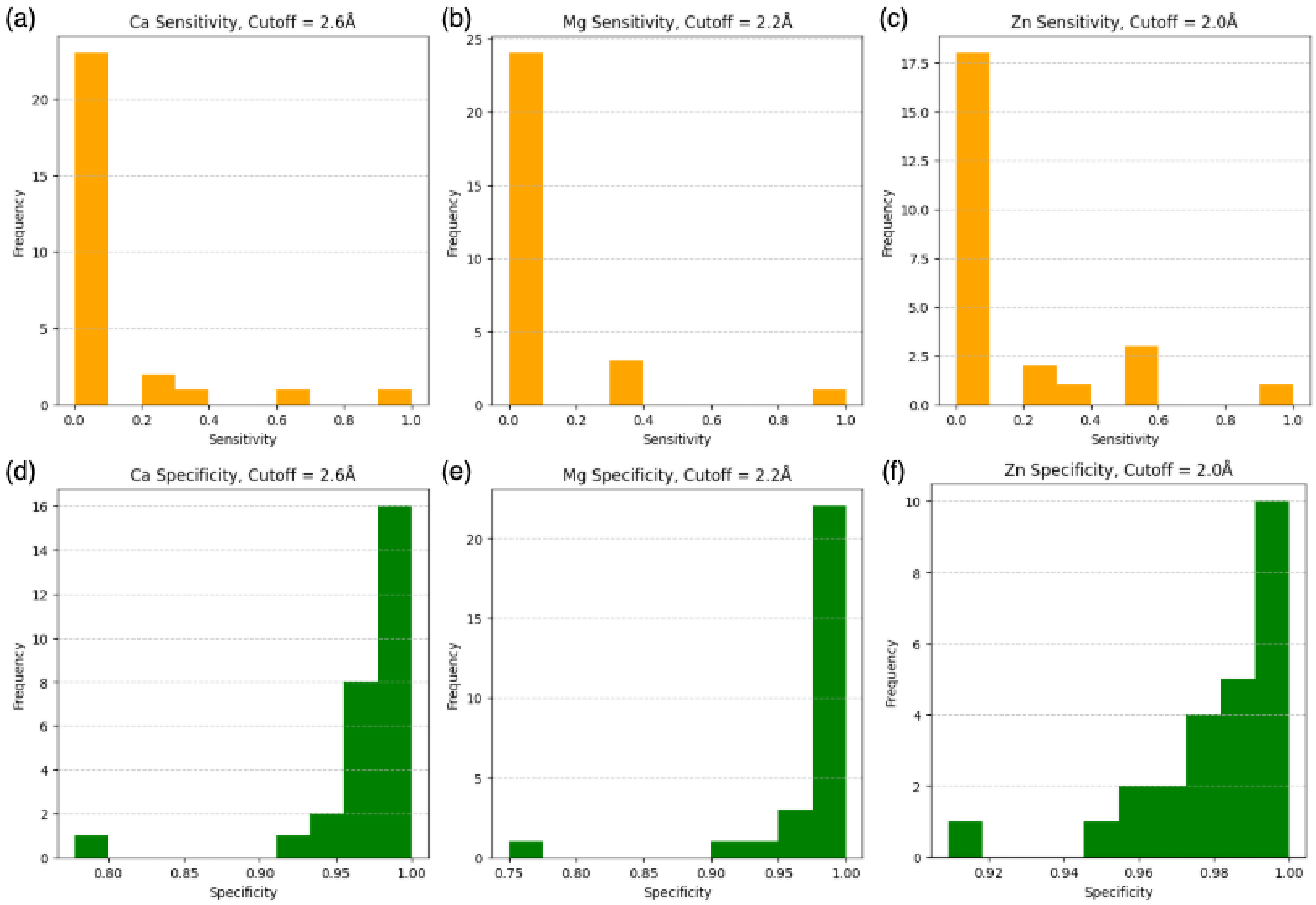
(a)
Ca Sensitivity, Cutoff = 2.6Å
Frequency
Sensitivity
(b)
Mg Sensitivity, Cutoff = 2.2Å
Frequency
Sensitivity
(c)
Zn Sensitivity, Cutoff = 2.0Å
Frequency
Sensitivity
(d)
Ca Specificity, Cutoff = 2.6Å
Frequency
Specificity
(e)
Mg Specificity, Cutoff = 2.2Å
Frequency
Specificity
(f)
Zn Specificity, Cutoff = 2.0Å
Frequency
Specificity

**Fig. S3**

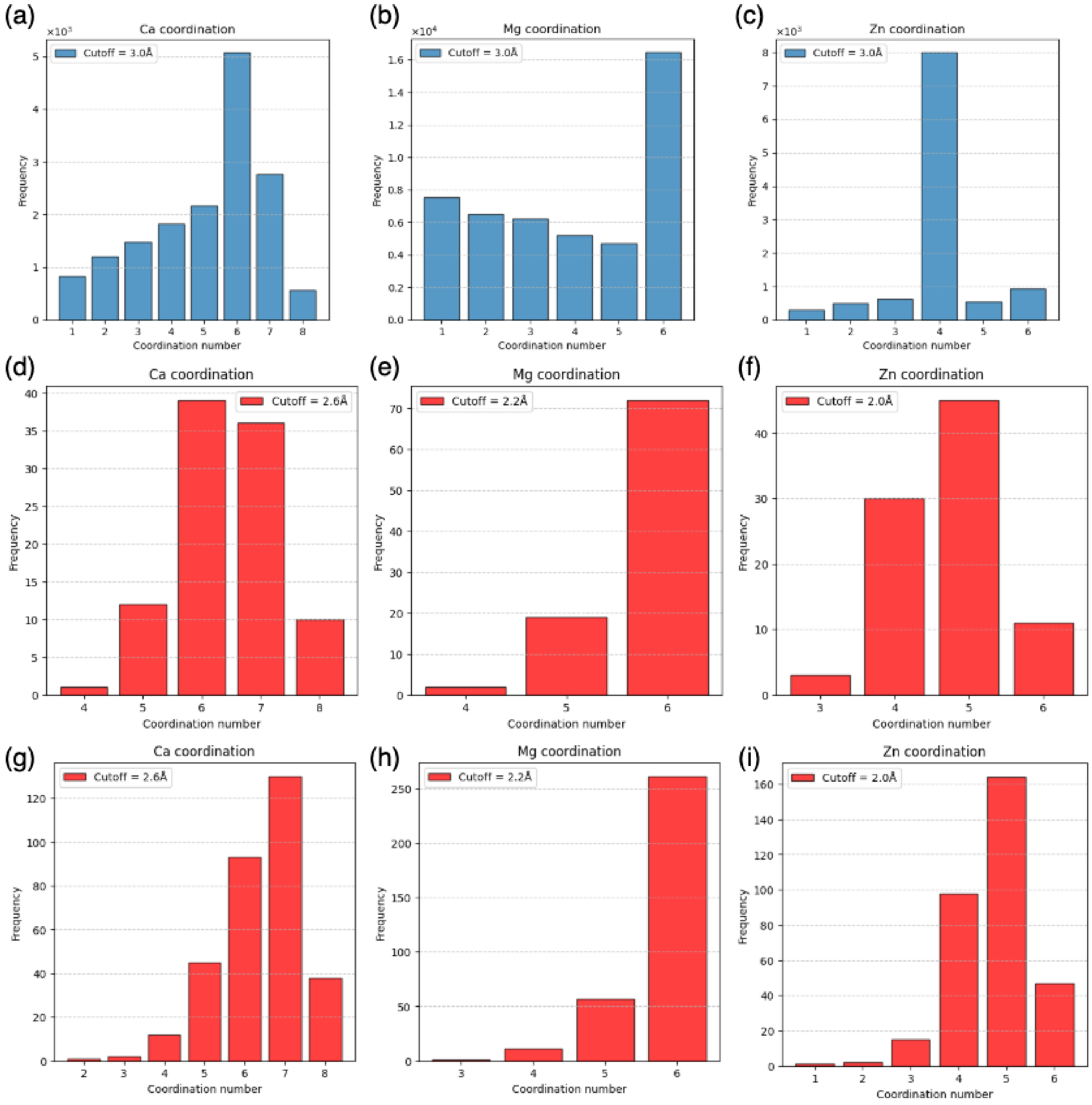
(a)
Ca coordination
Cutoff = 3.0Å
Frequency
Coordination number
(b)
Mg coordination
Cutoff = 3.0Å
Frequency
Coordination number
(c)
Zn coordination
Cutoff = 3.0Å
Frequency
Coordination number
(d)
Ca coordination
Cutoff = 2.6Å
Frequency
Coordination number
(e)
Mg coordination
Cutoff = 2.2Å
Frequency
Coordination number
(f)
Zn coordination
Cutoff = 2.0Å
Frequency
Coordination number
(g)
Ca coordination
Cutoff = 2.6Å
Frequency
Coordination number
(h)
Mg coordination
Cutoff = 2.2Å
Frequency
Coordination number
(i)
Zn coordination
Cutoff = 2.0Å
Frequency
Coordination number

**Fig. S4**
